# Calpains are required for efficient microtubule detyrosination

**DOI:** 10.1101/2021.07.08.451629

**Authors:** Julia Bär, Yannes Popp, Tomas Koudelka, Andreas Tholey, Marina Mikhaylova

**Affiliations:** AG Optobiology, Institute of Biology, Humboldt Universität zu Berlin, Berlin, Germany; Guest Group “Neuronal Protein Transport”, Center for Molecular Neurobiology, ZMNH, University Medical Center Hamburg-Eppendorf, Hamburg, Germany; Systematic Proteome Research & Bioanalytics, Institute of Experimental Medicine, Christian-Albrechts-University, Kiel, Germany

**Keywords:** Microtubules, detyrosination, vasohibin, calpain, mass spectrometry

## Abstract

Detyrosination is a major post-translational modification of microtubules (MT), which has significant impact on MT function in cell division, differentiation, growth, migration, polarity, and intracellular trafficking. Detyrosination of α-tubulin occurs via the recently identified complex of vasohibin 1/2 (vash1/2) and small vasohibin binding protein (SVBP). However, there is still remaining detyrosinating activity in the absence of vash1/2/SVBP, and little is known about the regulation of detyrosination. Using cellular and cell-free assays we showed that the calcium-dependent proteases calpains 1 and 2 regulate MT detyrosination. We identified new calpain cleavage sites in the N-terminal disordered region of vash1 using *in vitro* proteolysis followed by mass spectrometry. However, this cleavage did not affect the detyrosination activity of vasohibin. In conclusion, the regulation of MT detyrosination by calpains occurs via another yet unknown tubulin carboxypeptidase. Importantly, calpains’ calcium dependency could allow a fine regulation of MT detyrosination. Thus, identifying the calpain-regulated pathway of MT detyrosination can be of major importance for several basic and clinical research and should be focused on in future studies.

**Summary Statement:** The conventional calpains 1 and 2 play an important role in the regulation of microtubule detyrosination in a vasohibin independent way. Thus, they possibly control another still unknown tubulin carboxypeptidase.

## Introduction

In eukaryotic cells actin filaments, intermediate filaments, and microtubules (MT) are key components of the cytoskeleton. MTs play a crucial role in cell division, differentiation, growth, migration, polarity, and intracellular trafficking. Their structure, stability and functions are tightly regulated. MTs are hollow tubes with a well-defined polarity formed through head-to-tail polymerization of α- and β-tubulin heterodimers. They are about 24 nm in diameter and mainly build from 13 protofilaments. Polymerization of guanosine-5’-triphosphate (GTP)-bound tubulin-dimers leads to growth of MTs. Over time, β-tubulin hydrolyses GTP to guanosine-5’-diphosphate (GDP), which makes MTs more prone to depolymerization. A cap of GTP-bound tubulin will form at the plus end and protect the MT from disassembly. When hydrolysis catches up with polymerization, the tip will become GDP-bound, and the MT will depolymerize in an event called catastrophe. Upon reaching a GTP island or assembly of new GTP-bound tubulin, the MT will undergo a so-called rescue and will grow again. This continuous switch is called dynamic instability, as the MT overall persists, but its subunit composition changes (Dimitrov et al., 2008; Théry and Blanchoin, 2021). The more dynamic and fast-growing part of the MT is called plus end, whereas the more stable is the minus end (Desai and Mitchison, 1997). This intrinsic asymmetry is recognized by kinesins and dynein motor proteins that move along MTs in a directed fashion and mediate active long-distance cargo transport.

The tubulin family has expanded during evolution with expression of several α- and β-tubulin isoforms (8 and 9 in human, respectively). Both isotypes can undergo different post-translational modifications (PTMs), which may act as traffic signals and regulate stability and function of MTs, thus defining the so-called tubulin code (Garnham and Roll-Mecak, 2012; Janke, 2014). PTMs can label a subset of MTs as tracks for specific cargo-motor protein complexes, increase interaction with certain microtubule-associated proteins (MAPs), that can bundle MTs together, or change their stability by recruiting MT-severing proteins or polymerization promoting factors (Janke, 2014).

The removal of the C-terminal tyrosine of α-tubulin, called detyrosination, is one of the first discovered PTMs of MTs (Janke, 2014). Tubulin detyrosination occurs in a great variety of species ranging from invertebrates to humans and affects almost all α-tubulin isoforms (Janke, 2014; Redeker, 2010). In contrast to classical proteolytic cleavage, tubulin detyrosination is reversible. Upon depolymerization of MTs, the tyrosine residue can be re-ligated by tubulin tyrosine ligase (TTL), thereby returning αβ-tubulin dimers to the polymerization-depolymerization cycle. In fact, the presence or absence of the tyrosine residue has dramatic consequences for tumor progression, cardiomyocyte function and neuronal organization (Chen et al., 2018; Dubey et al., 2015; Heinz et al., 2017). Detyrosination is a very common modification in the mammalian brain and distortion of the tubulin turnover is observed in several pathological conditions. Thus, reduced MT stability has been reported in several neurodegenerative diseases including Alzheimer’s disease, Parkinson’s disease, and amyotrophic lateral sclerosis (Dubey et al., 2015; Yasuda et al., 2017). Tyrosination/ detyrosination of MT can act as a binary on/off switch for MT function (Garnham and Roll-Mecak, 2012). Detyrosinted (detyr-) MTs are more stable, protected from kinesin-13 induced depolymerization and serve as tracks for transport whereas tyrosinated (tyr-) MTs play a structural role (Kaul et al., 2014; Sirajuddin et al., 2014).

The identity of α-tubulin tyrosine carboxypeptidase (TTCP) was not known until recently when the vasohibin (vash)/small vasohibin binding protein (SVBP) complexes were discovered as the enzymes detyrosinating MTs (Aillaud et al., 2017; Nieuwenhuis et al., 2017). Vasohibins are represented by two similar and functionally redundant proteins: vash1 and its homolog vash2. They belong to the group of transglutaminase-like cysteine proteases but contain a non-canonical Cys-His-Ser catalytic triad (Aillaud et al., 2017). Vasohibins are widely distributed in eukaryotes, have broad tissue expression, and vash1 is more abundant, especially in the brain (Aillaud et al., 2017).

Since detyrosination has such a strong impact on MT functions, there are likely regulatory mechanisms, which can control the activity of vasohibins spatially and temporally. Both vash1 and vash2 have some basal TTCP activity, which is stronger in case of vash2 (van der Laan et al., 2019).The enzymatic activity of both vasohibins is facilitated by association with the small chaperone protein SVPB *in vivo* and *in vitro*. However, if and how vasohibin’s detyrosination activity is regulated besides via SVBP binding remains unknown. Interestingly, before their identification as TTCPs, vasohibins were mostly studied as secreted proteins without any classical signal peptide sequences. Their secretion required association with SVBP but the exact mechanisms of their extracellular release are not yet understood (Kadonosono et al., 2017). The calcium-dependent non-lysosomal cysteine protease calpain 1 was shown to cleave vash1 from the N- and C-terminus, thereby altering its angiogenic activity in the extracellular space (Saito et al., 2016). Enzymatic activity of the classical calpains 1 and 2 is controlled by calcium. Intracellular calcium concentration can rapidly change depending on the physiological status of a cell, which makes calpains an attractive target in search of the mechanism regulating detyrosination of MTs. Therefore, it is still an open question whether calpain-dependent proteolysis of vasohibins could be the mechanism controlling MT detyrosination. Furthermore, several human cell lines (e.g HEK293T, UO2S) depleted of vasohibin/SVBP or mice lacking SVBP still show some remaining detyrosinating activity suggesting that there might be other enzyme(s) responsible for this microtubule modification (Aillaud et al., 2017; Liao et al., 2019; Nieuwenhuis et al., 2017; Pagnamenta et al., 2019).

In this study, we show that proteolytic activity of calpain 1 and 2 is required for efficient MT detyrosination in HEK293T cells. This is not due to the regulation of vasohibin activity via cleavage and not via direct modification of MT by calpain, suggesting that calpain acts upstream of a still unidentified TTCP.

## Results

### Calpain inhibition reduces microtubule detyrosination in HEK293T cells

To narrow down the class of proteases involved in detyrosination of α-tubulin either via regulation of vasohibins or as additional TTCPs, we treated HEK293T cells with parthenolide (TTCP activity inhibitor), potato carboxypeptidase inhibitor (PCI, a general metalloproteinase inhibitor), or calpeptin (calpain 1 and 2 inhibitor). Interestingly, a 30 min treatment with 60 μM calpeptin leads to a strong decrease in the amount of detyr-MTs as evidenced by immunostainings or immunoblotting with a detyr-α-tubulin antibody (Fig. 1B, C). To differentiate between effects on MT detyrosination via a peptidase inhibition or a tyrosine ligation via TTL (Fig. 1A), we performed calpeptin treatments on HEK293T cells where the MTs were stabilized by taxol for 30 min. Since TTL acts on soluble tubulin dimers, taxol treatment would prevent MT depolymerization, thus limiting the pool of soluble tubulin. Taxol alone increased the amount of detyr-tubulin as seen by immunocytochemistry (ICC) and immunoblotting. This is in line with the notion that long-lived MTs accumulate detyr-tubulin (Garnham and Roll-Mecak, 2012). This effect was strongly reduced by co-application of calpeptin, which indicates that calpain activity is required for the proper function of TTCP (Fig. 1B, C).

**Figure 1.**
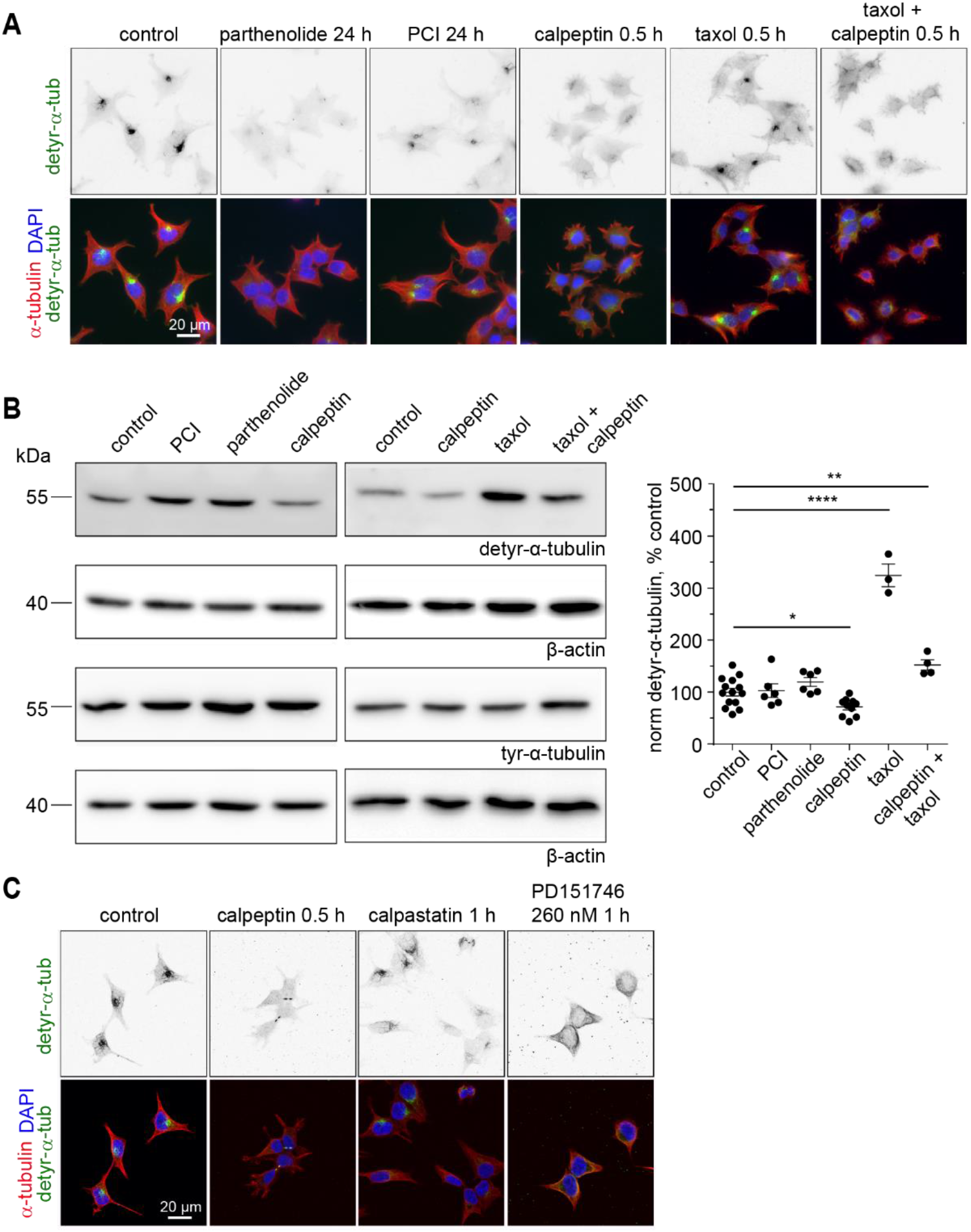
Microtubule detyrosination in HEK293T cells depends on classical calpains. A) Representative fluorescence widefield images of untreated HEK293T cells, and cells treated with indicated inhibitors and stained for detyrosinated (green) and total (red) tubulin. B) Treatment of cells with the calpain inhibitor calpeptin reduces the amount of detyrosinated MT. Example western blot and quantification. Data are represented as mean ±SEM. Dots represent individual measurements from 3 independent experiments. 1-Way-ANOVA (p<0.0001) with Dunnett‘s multiple comparison test against control. * p≈0.04, ** p≈0.004, **** p<0.0001. C) Representative maximum projections of fluorescence confocal images of HEK293T cells treated with DMSO as control and different calpain inhibitors show the importance of calpain in MT detyrosination. Note the strong reduction in detyr-tubulin staining intensity (green) upon calpeptin treatment.

To distinguish between different calpains that might be involved in the regulation of TTCP we performed similar experiments using calpastatin - the only absolutely specific inhibitor for classical calpains, and PD151746 - so far the most specific calpain-1 inhibitor with a 20x higher selectivity than for calpain-2 (Fig. 1C) (Ono et al., 2016). All three inhibitors had a similar effect on detyrosination of MTs pointing to the involvement of classical calpains on MT detyrosination.

To further verify the involvement of calpains, we next performed knockdown experiments with calpain 1 and 2 siRNAs (Fig. 2). The specificity and functionality of the knockdown in HEK293T cells were confirmed by western blot analysis after 24 h of transfection (Fig. 2A). A strong reduction in MT detyrosination upon calpain 1 and 2 knockdown was visible in fluorescent immunostainings already after 24 h (Fig. 2B). To validate these results, we performed a western blot analysis of HEK293T cells co-transfected with vash-1-GFP and respective calpain siRNAs (Fig. 2C). Co-transfection with either calpain 1 or 2 siRNAs led to a significant decrease in detyr-MTs. The combination of both siRNAs turned out to be less efficient, probably due to the reduced amount of the individual siRNAs (Fig. 2C, D). Taken together, these data indicate that conventional calpains play a role in the regulation of the MT cytoskeleton and calpain activity is required for efficient MT detyrosination.

**Figure 2.**
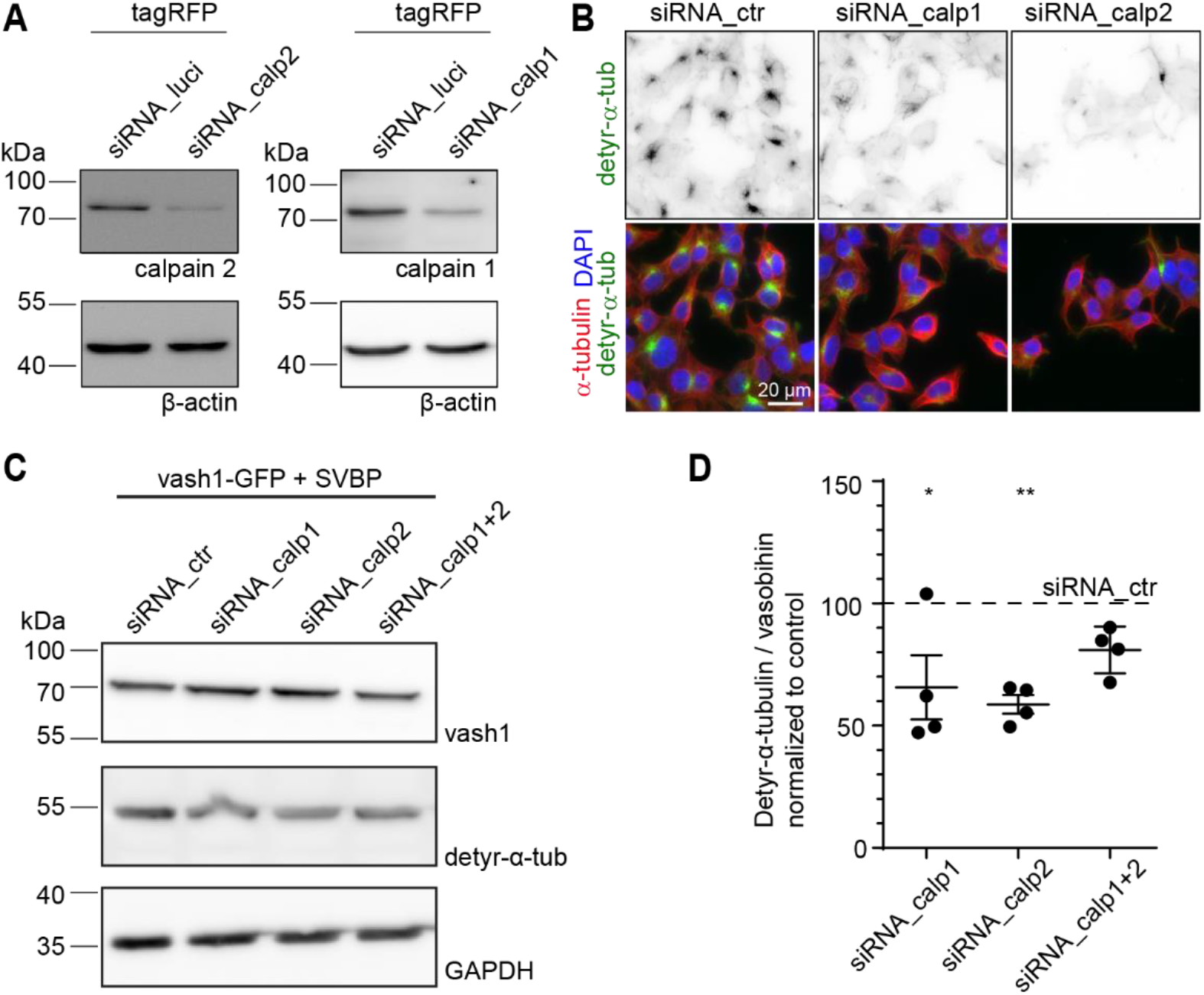
Knockdown of calpain 1/2 leads to reduction of detyrosinated α-tubulin. A) Confirmation of knockdown of calpain1/2 in HEK293T using western blotting. HEK293T cells were transfected with tagRFP and respective siRNAs for 24 h. β-actin was used as a loading control. B) Representative maximum projections of fluorescence widefield images of HEK293T cells transfected with different siRNAs for 24 h and stained for detyr-tubulin (green) and total tubulin (red) shows reduction of detyrosination. C) HEK293T cells were double transfected with vash1-GFP-SVBP and siRNA for calpain as indicated for 24 h. Cells were harvested, lysed, and subjected to western blotting. D) Quantification of C) n= 4 independent experiments with 1-3 samples per group. 1-Way-ANOVA [F(3,12)=6.3 p=0.008] with Dunnett’s post-hoc test. * p<0.05, ** p<0.01.

### Calpain inhibition does not affect MT growth in HEK293T cells

As MT detyrosination and stability strongly correlate (Garnham and Roll-Mecak, 2012) we next assessed the MT dynamics in HEK293T cells upon calpain knockdown (Fig. 3). HEK293T cells were co-transfected with the MT plus end marker EB3-GFP and the respective siRNAs for calpain 1 and 2. Using live total internal reflection fluorescence imaging and tracking of EB3 comets (Fig. 3A) we could not find changes in EB3 comet number (Fig. 3B), speed, or track lengths upon knockdown of calpain (Fig. 3C). Although MT detyrosination is strongly reduced upon calpain KD, it does not seem to affect MT dynamics.

**Figure 3.**
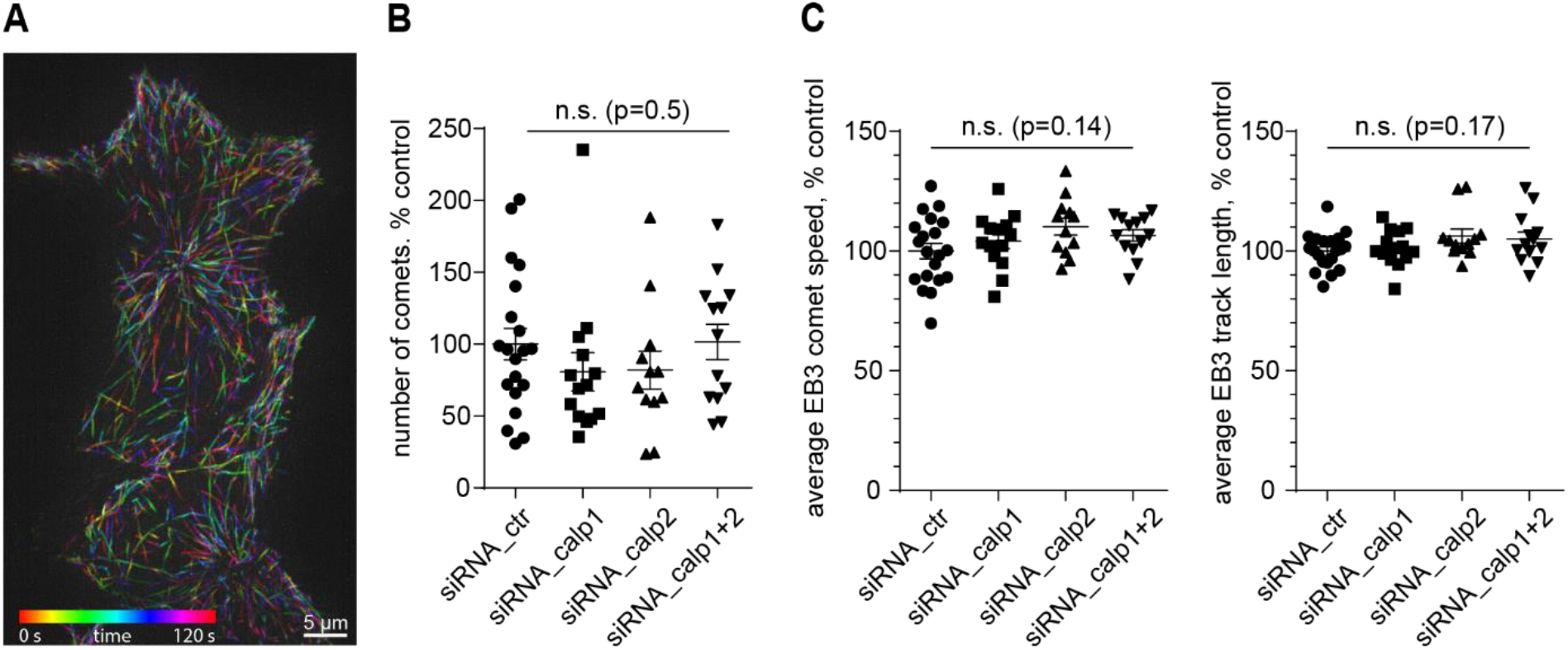
Knockdown of calpain 1 or 2 has no effect on MT growth/dynamics. A) Representative image of EB3 comets imaged over 2 min in HEK293T cells. Temporal color code shown in bottom left. B) Number of EB3-comets per cell area is unchanged upon knockdown of calpains. 1-Way ANOVA. n= 20 (siRNA_ctr), 14 (siRNA_calp1), 12 (siRNA_calp2), 13 (siRNA_calp1+2) cells from 2 independent experiments. Data are represented as mean ±SEM. C) Quantification of comet speed and track length. 1-Way ANOVA. n= 20 (siRNA_ctr), 14 (siRNA_calp1), 12 (siRNA_calp2), 13 (siRNA_calp1+2) cells from 2 independent experiments. Data are represented as mean ±SEM.

### Effect of vasohibin cleavage by calpain 1 on MT detyrosinating activity

Calpain 1 and 2 are the conventional and ubiquitously expressed calpain family members. They are heterodimers consisting of an identical small regulatory subunit CAPNS1 of 28 kDa size and a distinct catalytic 80 kDa subunit, CAPN1, or 2, respectively. Calpain 1 and 2 have a broad peptidase activity without clear sequence specificity (Shinkai-Ouchi et al., 2016). Predictions for cleavage sites based on amino acid sequence using machine learning indicate that calpains require longer stretches (P14-P10’ and P20 to P20’ for calpain 1 and 2, respectively) for substrate cleavage (Sorimachi et al., 2012).

Thus, it is unlikely that calpains are detyrosinating α-tubulin themself. It is more likely to be upstream of TTCP. Vash1 was previously proposed to be a substrate of calpain 1 (Saito et al., 2016) and potential cleavages have been published in context of its anti-angiogenetic activity and tumor microenvironments: active 44 kDa full-length vash1 is released and proteolytically processed at the N- and C-terminus resulting in active 36 kDa and inactive 27 kDa products respectively (Saito et al., 2016; Sonoda et al., 2006). However, the consequences of vasohibin cleavage on its cytosolic functions, specifically detyrosination of MT, are completely unknown. Therefore, we next performed a set of cell free assays to investigate if calpain 1 is indeed cleaving vash1, and if it is upstream of vasohibin in the regulation of MT detyrosination (Fig. 4, 5).

**Figure 4.**
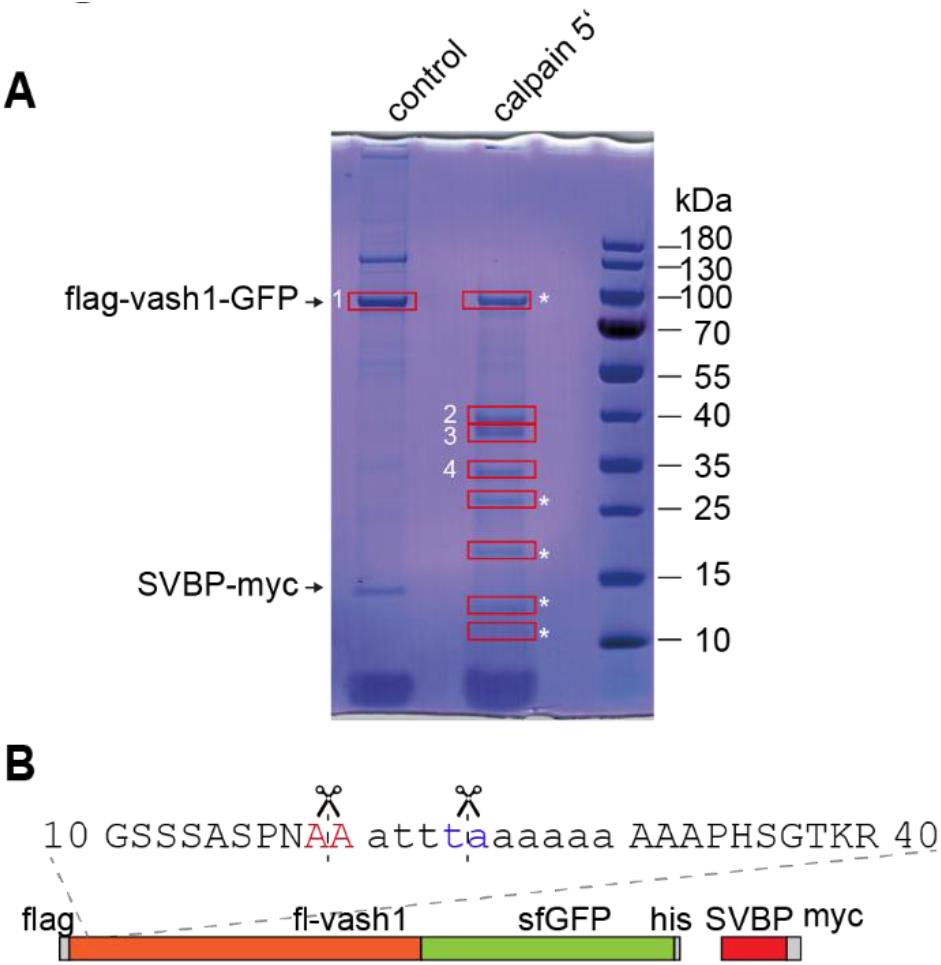
Identification of calpain cleavage sites within vash1 by mass spectrometry A) SDS-PAGE showing purified SVBP-myc and co-purified vash1-GFP with and without on bead-calpain digestion (>20 U, 5 min). Boxed bands were cut and used for mass spectrometry. Asterisks indicate bands enriched in calpain 1 catalytic subunit. B) Identified N-termini upon calpain cleavage after trypsin and chymotrypsin digest and N-terminal dimethylation. Highlighted are newly identified N-termini present in both trypsin and chymotrypsin digest. Small letter residues are contained in mouse vash1, but not human. Also see Tables S1 and S2.

**Figure 5.**
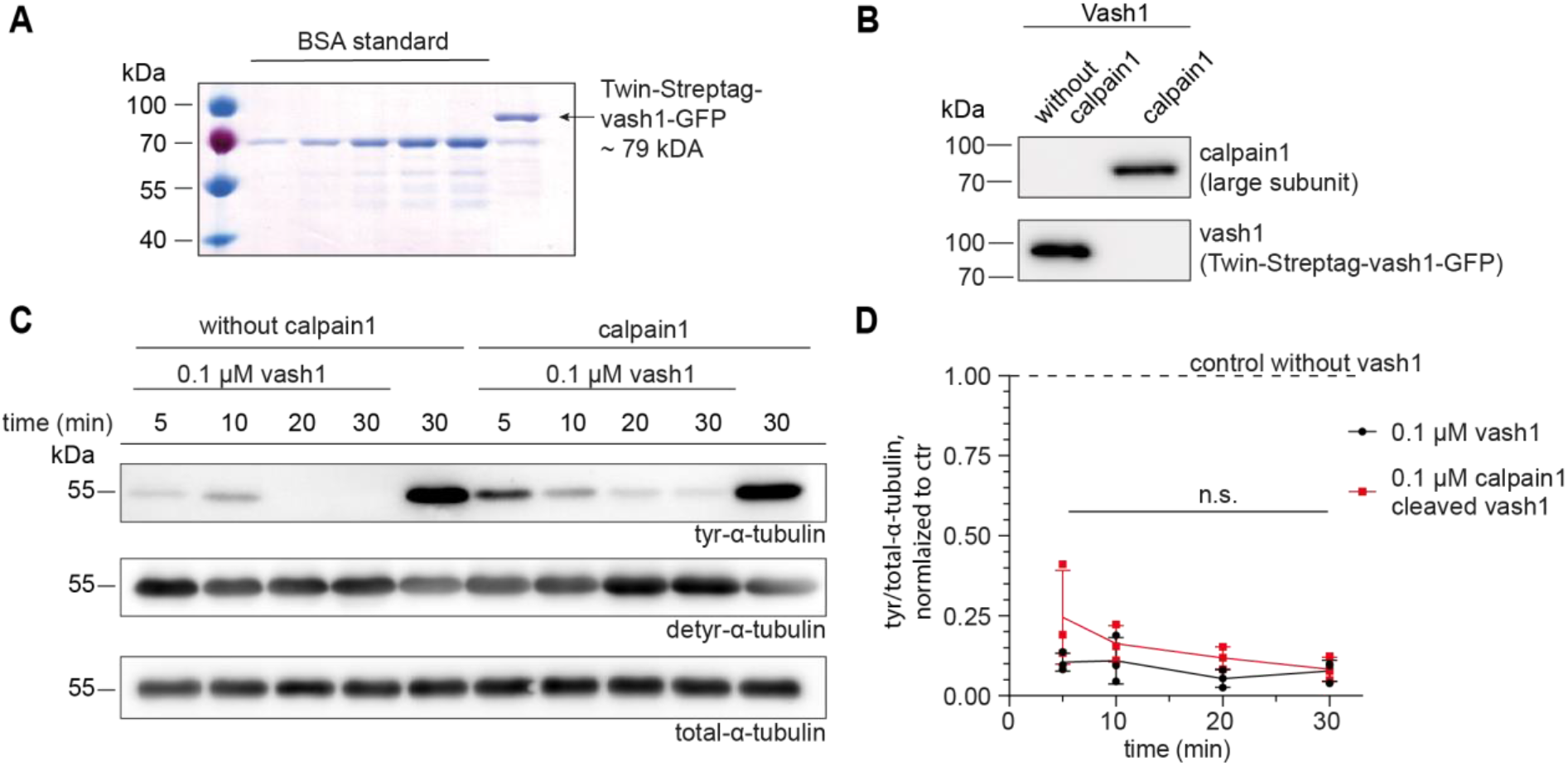
Calpain 1 cleavage has no influence on vash1-mediated MT detyrosination *in vitro*. A) Coomassie-stained 10 % SDS-polyacrylamide gel of purified twin-Streptag-vash1-GFP from HEK293T cells. Vash1: 1.5 μg in 5 μL, BSA standard: 0.25, 0.5, 1.0, 1.5, 2.0 μg. B) Western blot showing cleavage of purified vash1 in presence of calpain 1. C) Purified vash1 causes strong loss of tyr-α-tubulin signal and increase in detyr-α-tubulin signal in western blot from MTs *in vitro*, independent of prior cleavage by calpain 1 and incubation time. D) Quantification of tyr/total-α-tubulin ratio normalized to a negative control without vash1. n=3 independent experiments. 2-Way ANOVA followed by Tukey’s multiple comparisons test (condition ± calpain1 p=0.1701).

To isolate vash1 from HEK293T cells independent of N- or C-terminal cleavage, we decided to co-purify it with SVBP-myc as a bait, as the SVBP binding side lies within the central region of vash1 (aa 133-144 and 274-282) (Kadonosono et al., 2017). After on-bead calpain digestion using purified recombinant calpain 1 and subsequent separation on an SDS gel, prominent bands were excised and used to identify potential new N-and C-termini termini (Fig. 4A). This was achieved by using dimethyl labelling at the protein level followed by enzymatic digestion in the presence 50 % heavy water (H_2_^18^O) and LC-MS analysis (Deng et al., 2015; Schnölzer et al., 1996). While new C-termini could only be identified within the linker-region and GFP (not shown), several new N-termini were found within the alanine-rich N-terminus of vash1 (Fig. 4B, Table S1-S2). Namely between residues 19A.20A and 25A.26A, which were identified in both the chymotrypsin and trypsin experiments and with a large number of peptide spectral matches (PSMs). These positions are in agreement with the cleavage specificity of calpain 1 according to the MEROPS peptidase database (https://www.ebi.ac.uk/merops/) (Rawlings et al., 2017) in which calpain prefers alanine in the P1’ site. Another less abundant cleavage site, S14.A15, was also identified in digestions experiments. This site contains an alanine residue in the P1’ position and a proline residue in the P3’ position, which is in line with calpain’s cleavage site specificity (Figure S1). Moreover, recently published structures of vasohibins suggest a disordered N- and C-terminus terminus (Adamopoulos et al., 2019; Li et al., 2019; Liao et al., 2019; Wang et al., 2019) and therefore the alanine-rich N-terminus would be accessible to calpain, which prefers interdomain unstructured regions in its substrate binding pocket (Hosfield et al., 1999; Moldoveanu et al., 2002; Sorimachi et al., 2012). Of note, previously reported cleavage corresponding to amino acids 86/87 and 328/329 in the mouse protein (Sonoda et al., 2006) could not be confirmed.

After identifying these new cleavage sites, we investigated the functionality of cleaved vash1 on MT detyrosination by performing a cell free MT assay (Fig. 5). To obtain high purity, soluble, eukaryotically produced vash1, the vash1 coding sequence was cloned into a twin-strep-tag together with SVBP and purified from HEK293T cells using strep-Tactin beads (Fig. 5A). Tyrosinated MTs were prepared from porcine brain tubulin by 2 rounds of polymerization/depolymerization and incubated with purified twin-strep-tag-vash1-GFP, with or without prior calpain 1 digestion, for different time intervals from 5 to 30 min (Fig. 5B, C). Western blotting analysis confirmed that the presence of calpain 1 leads to cleavage of vash1, as the full-length-vash1 band was no longer detectable (Fig. 5B). As expected the addition of vash1 lead to a strong reduction in tyrosinated α-tubulin, and an increase in detyrosinated α-tubulin (Fig. 5C, D). However, this effect was largely independent of the presence of calpain 1, indicating that calpain 1 cleavage of vash1 does not affect its MT detyrosinating abilities (Fig. 5C, D). This experiment can also rule out the possibility that calpain 1 itself cleaves the C-terminal tyrosine of α-tubulin.

To further confirm these findings in a cellular context, we cloned and overexpressed different vash1 truncation constructs in HEK293T cells and quantified MT detyrosination after immunostaining: 87-375-vash1 and 87-328-vash1, representing the previously described ΔN and ΔNΔC (Sonoda et al., 2006), and the newly identified 20/25-375 (Fig. 6A). While the full-length vash1 expectedly lead to a strong increase in MT detyrosination, 87-375-vash1 and 87-328-vash1 did not have an effect, indicating that N- and C-terminal truncation renders the protein functionally inactive in the context of MT detyrosination (Fig. 6B, C). The newly identified cleavage of calpain 1 in the N-terminus of vash1 (20/25-375) leads to the same increase in MT detyrosination as the full-length construct (Fig. 6B, C), confirming that calpain 1 does not regulate vasohibin’s function in the MT detyrosination pathway, opening a possibility that calpains act upstream of a yet unknown TTCP.

**Figure 6.**
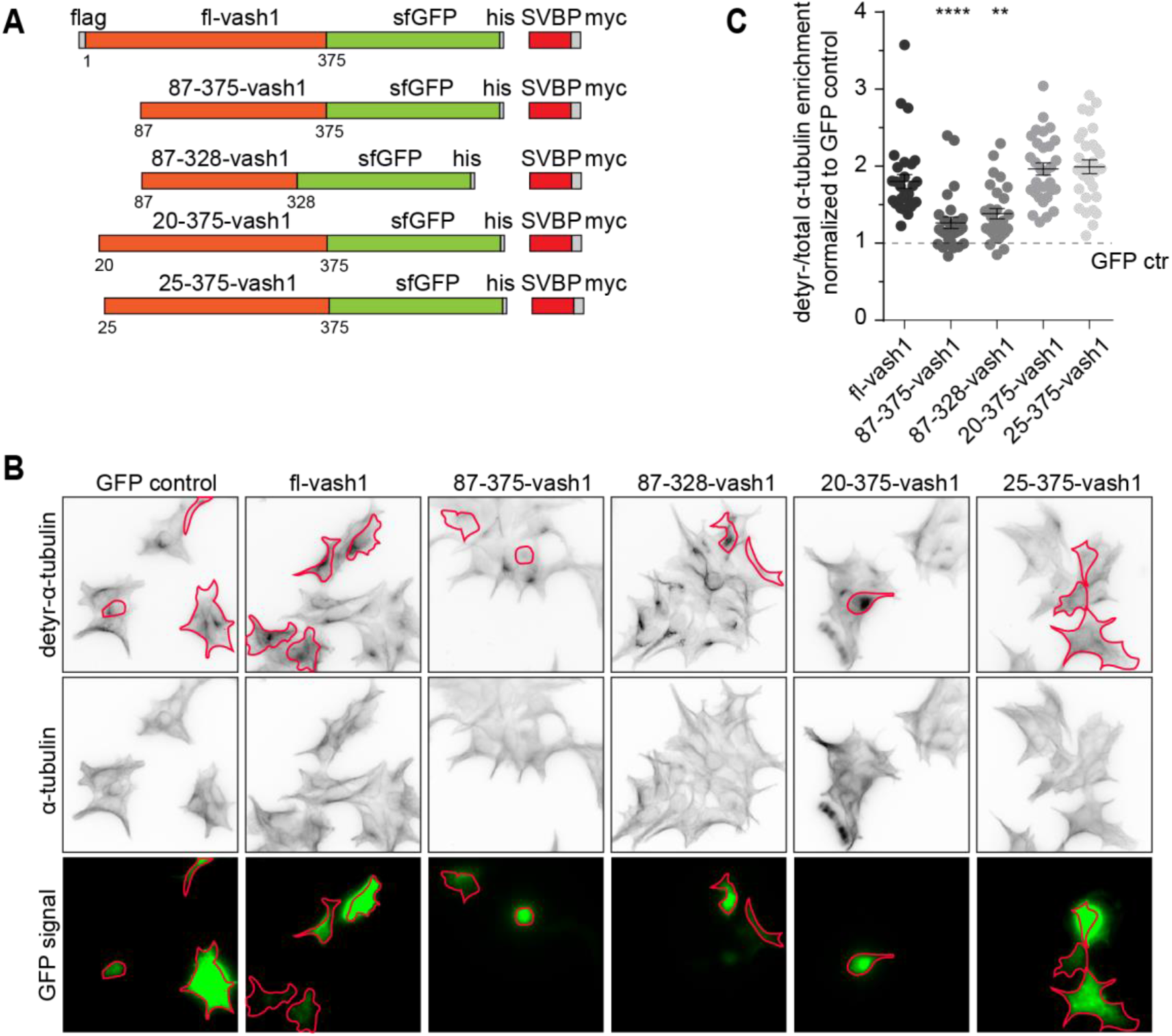
Effects of vash1 truncation constructs on MT detyrosination. Overexpression of full-length vash1 and calpain-cleaved vash1 leads to MT detyrosination in HEK293T cells. A) Scheme showing vash1 and its truncation constructs used for overexpression in HEK293T cells. A) Representative widefield fluorescence images of HEK293T cells transfected with the indicated constructs and stained for detyr-tubulin and total α-tubulin. Transfected cells are outlined in green. B) Quantification of detyr/total tubulin ratio in transfected HEK293T cells compared to untransfected cells, normalized to GFP control. n= 30 (GFP), 30 (fl-vash1), 27 (87-375-vash1), 28 (87-328-vash1), 30 (20-375-vash1), 29 (25-375-vash1) in 3 independent experiments. Kruskal-Wallis test (p<0.0001) with Dunn’s multiple comparison test. **** p<0.0001, ** p=0.0035.

## Discussion

In this work, using an inhibitor screen and immunostainings in HEK293T cells we discovered that the conventional calpains 1 and 2 are involved in the regulation of MT detyrosination. Specifically, pharmacological inhibition of calpain 1 or 2 and siRNA knockdown approaches resulted in a strong decrease of detyrosinated MT. MT stabilization by application of taxol confirmed a decrease in detyrosination, i.e., TTCP activity, rather than an increase in TTL activity. Despite the strong changes in MT detyrosination, MT number and growth seems largely unchanged upon calpain knockdown suggesting that calpains act on specific aspects of the MT cycle. Although, one should keep in mind that the used method might not be sensitive enough to detect changes in very short comet tracks. MT polymerization rates are well analyzable and unchanged upon calpain knockdown. Therefore, future studies should address the effects of calpains on MT stability, for instance by examining catastrophes and rescues.

The profound effect of calpain on MT detyrosination could either be explained by regulation of the TTCP vash1/2/SVBP complex activity by calpains or by the existence of another detyrosination pathway which is sensitive to calpain. Since vash1 was proposed to be a calpain 1 substrate (Saito et al., 2016), we set out to examine this possibility by performing *in vitro* cleavage assays followed by mass spectrometry analysis to identify newly created N-termini. We found that calpain 1 cleaves mouse vash1 (NP_796328.2) in an alanine-rich stretch in the N-terminus. However, this cleavage did not affect the activity of vash1 on microtubule detyrosination, as we could show in cell free assays and in HEK293T cells. Moreover, we could not confirm the previously described cleavage products 87-375-vash1 and 87-328-vash1, corresponding to the N-terminally cleaved 36 kDa (ΔN) and the N- and C-terminally cleaved 27 kDa (ΔNΔC) form of human vash1 from (Sonoda et al., 2006). The authors identified these cleavage products in cell culture media collected from endothelial cells, co-cultured with various cancer cell lines, and showed changed angiogenic properties of vash1. While ΔN is still active, ΔNΔC is rendered inactive (Saito et al., 2016). Based on the assumption that limited proteolytic cleavage requires basic residues (Seidah and Chrétien, 1999) both truncation variants were identified in a mutagenesis screen where arginines were mutated to alanines. Calpains are calcium-dependent proteases with a broad endopeptidase activity without clear sequence specificity (Shinkai-Ouchi et al., 2016), but calpain 1 seems to prefer alanine in the P1’ site (MEROPS peptidase database) (Rawlings et al., 2017). Both cleavage sites for ΔN and ΔNΔC however do not show this alanine site. This might be a reason, why we could not identify those previously described cleavage products. Additionally, we used mouse vash1, while studies on the angiogenetic effects of vash1 are based on the human protein. Both share a 93 % sequence identity, but the mouse sequence contains 10 additional amino acids in the N terminus (aa 21-30, Fig. 4B), a stretch that is highly enriched in alanines, and contains the newly identified calpain cleavage sites.

The shorter cleavage products 87-375-vash1 and 87-328-vash1 were ineffective in increasing MT detyrosination in HEK293T cells. We conclude, that the cytosolic (e.g. detyrosination) activity of vash1 is independent of its angiogenic extracellular effect and both are regulated independently. Vash1 seem to be present ubiquitously in the brain (http://www.proteinatlas.org) (Uhlén et al., 2015), but at low to medium levels (and is currently not possible to detect in ICCs (unpublished data, personal communication Marie-Jo Moutin), while vash2 expression might be even lower (http://www.proteinatlas.org) (Uhlén et al., 2015). However, both have a strong effect on MT detyrosination, suggesting that low proteins amounts are sufficient for proper cellular functioning and possibly pointing to increased importance of regulation of vasohibin activity.

Calpain 1 and 2 as major non-lysosomal proteases play an important role in cell signaling due to the limited proteolysis of their substrates - both in physiological and pathological conditions (Moldoveanu et al., 2002). Proper neuronal branching, spine complexity and therefore hippocampal long-term plasticity and spatial memory depend on the presence of calpains (Amini et al., 2013; Zadran et al., 2010). Excessive calcium influx, occurring for instance during stroke, leads to increased calpain activity and MT destabilization, cleavage of MAP2, spectrin, internexin and others, finally leading to the breakdown of the cytoskeleton and cell death. The conventional calpains 1 and 2 seem to affect a still unknown TTCP rather than regulating vasohibin activity. This could provide another explanation why SVBP knockout mice show only an approx. 40 % reduction in detyr-MT (Pagnamenta et al., 2019) that might not fully be accounted for by the presence of the C-terminal tyrosine-lacking α-4-tubulin. Considering that inhibition of calpains leads to a loss of MT detyrosination it seems that calpain is an activator of this unknown TTCP. Regulation of PTMs of MTs and calpains has been shown to be important for several cellular processes in various cell types. The detyrosination status of MTs is essential for the function of the mitotic spindle, formed during cell division. Inhibition of detyrosination leads to misalignment of pole-proximal chromosomes during chromosome congression and delayed mitotic progression. This process depends on CENP/E (kinesin 7), which strongly prefers detyrosinated MTs (Barisic et al., 2015). During cell division, calpain 2 is upregulated in the G2-M phase and a reduction or inhibition of calpain 2 causes chromosome misalignment due to polar ejection force impairment (Honda et al., 2004). Thus, both (PTM and calpain activity) play together in maintaining proper cellular function.

Calpains calcium dependency can give rise to a fine regulation of MT detyrosination. This could be especially important in excitable cells, such as muscle cells and neurons, that underly strong changes in intracellular calcium levels and therefore calpain activity. In neurons, MTs play a major role in intracellular transport. Using motor-PAINT (motor protein-based point accumulation for imaging in nanoscale topology) and expansion microscopy it was recently shown that neurons contain bundles of spatially segregated stable and dynamic MTs with respective post-translational modifications (Jurriens et al., 2021; Tas et al., 2017). As both types of MTs are oriented in opposing directions and are preferred by different motor proteins, this allows for selective entry of motor proteins into either dendrites or axons. Therefore, PTMs of MTs, such as detyrosination, play a major role in the regulation of intracellular transport and neuronal morphology. Additionally, it was shown by immunostainings and fluorescent peptidase activity readouts in combination with calpain inhibitors, that calpain 1 and even more notably calpain 2, are present and active in neurites of pyramidal neurons *in vitro* and *in vivo* (Mingorance-Le Meur and O’Connor, 2009). This seems to be most prominent in axons and apical dendrites, the places with most microtubules detyrosination within neurons. The importance of MT detyrosination is further supported by the recent discovery of several families carrying SVBP “null” mutations, that lead to severe neurological deficits (Iqbal et al., 2019; Pagnamenta et al., 2019). Thus, identifying the calpain-regulated pathway of MT detyrosination can be of major importance for several basic and clinical research and should be focused on in future studies.

## Methods

### Cloning

All used plasmids and cloned construct are listed in table 1. For cloning of the vash1 truncation constructs, respective nucleotides were amplified via PCR from the full-length construct with primers containing restriction sites XbaI and AgeI and cloned back into the full-length backbone via standard restriction digestion & ligation cloning protocols. vash1-87-375-sfGFP-his IRES SVBP-myc and vash1-87-328-sfGFP-his IRES SVBP-myc correspond to the N-terminally cleaved 36 kDa and the N-and C-terminally cleaved 27kDa form of human Vasohin 1 from (Sonoda et al., 2006). The Twin-strep-tag-vash1-GFP IRES SVBP construct was obtained by cold fusion cloning of a gBlock containing the twin-strep-tag (TwinStrep-mCherry empty vector was a gift from A. Aher and A. Akhmanova, Utrecht University) into the Bsu15I digested full-length vash construct.

**Table 1:**
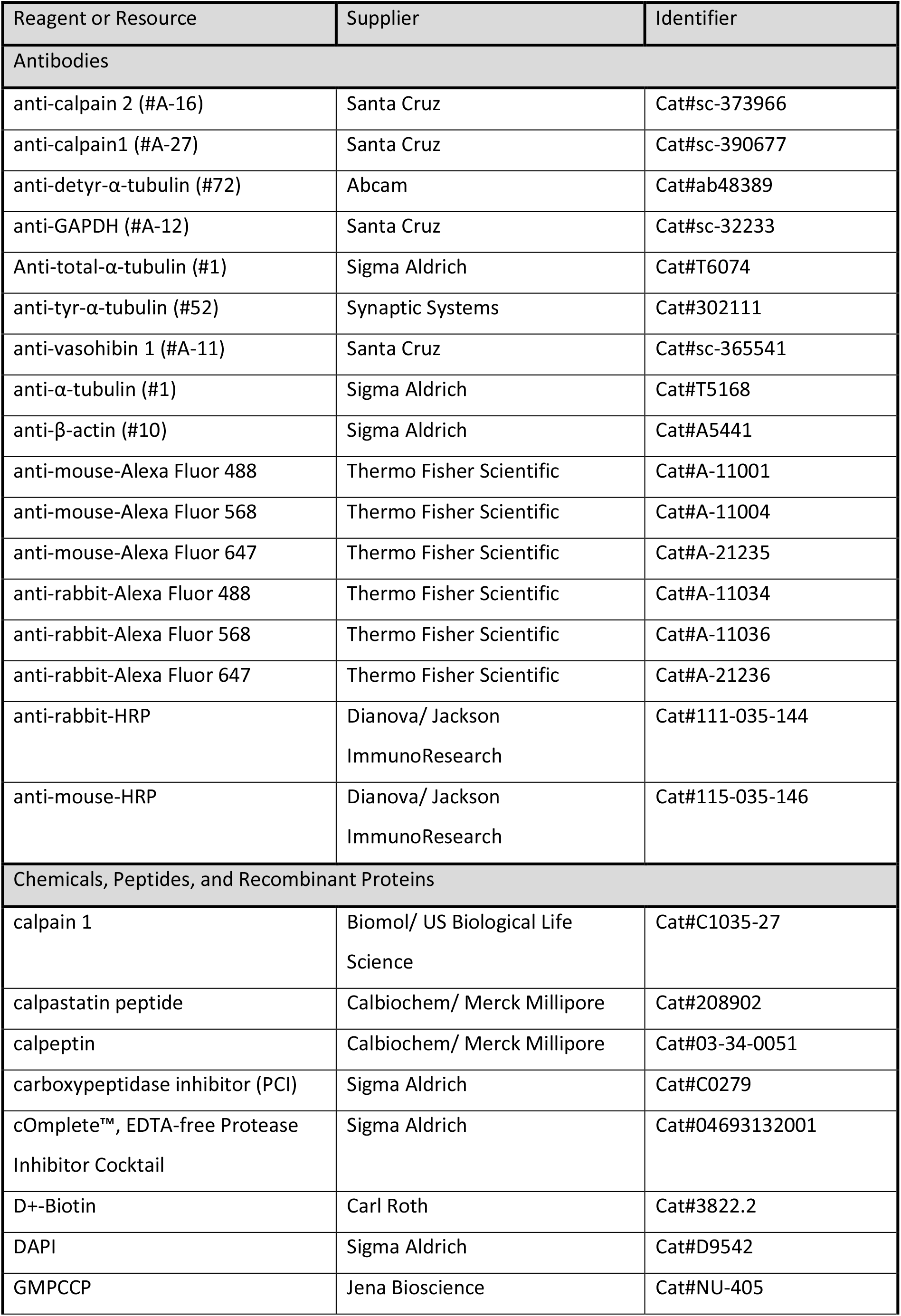
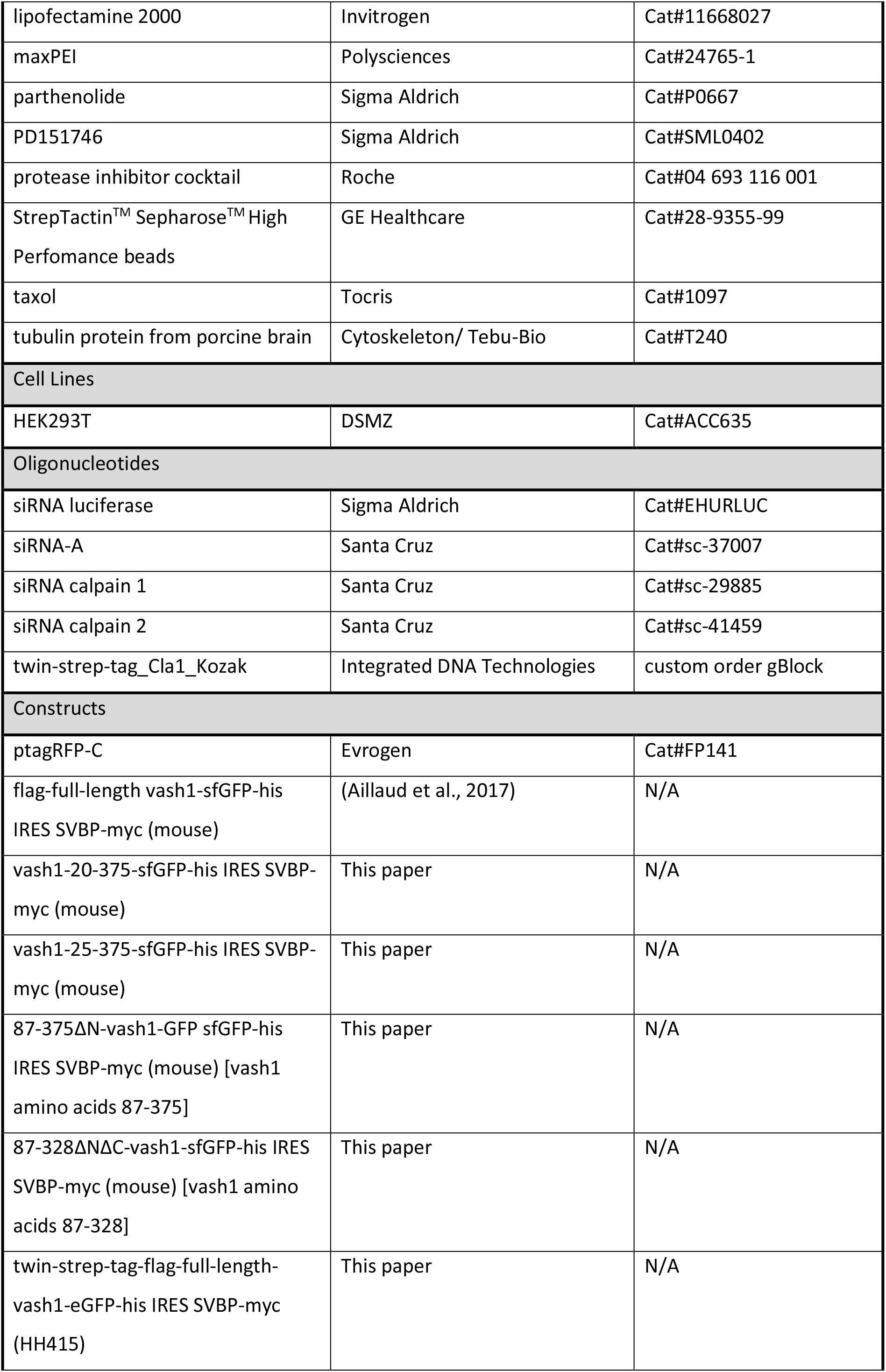
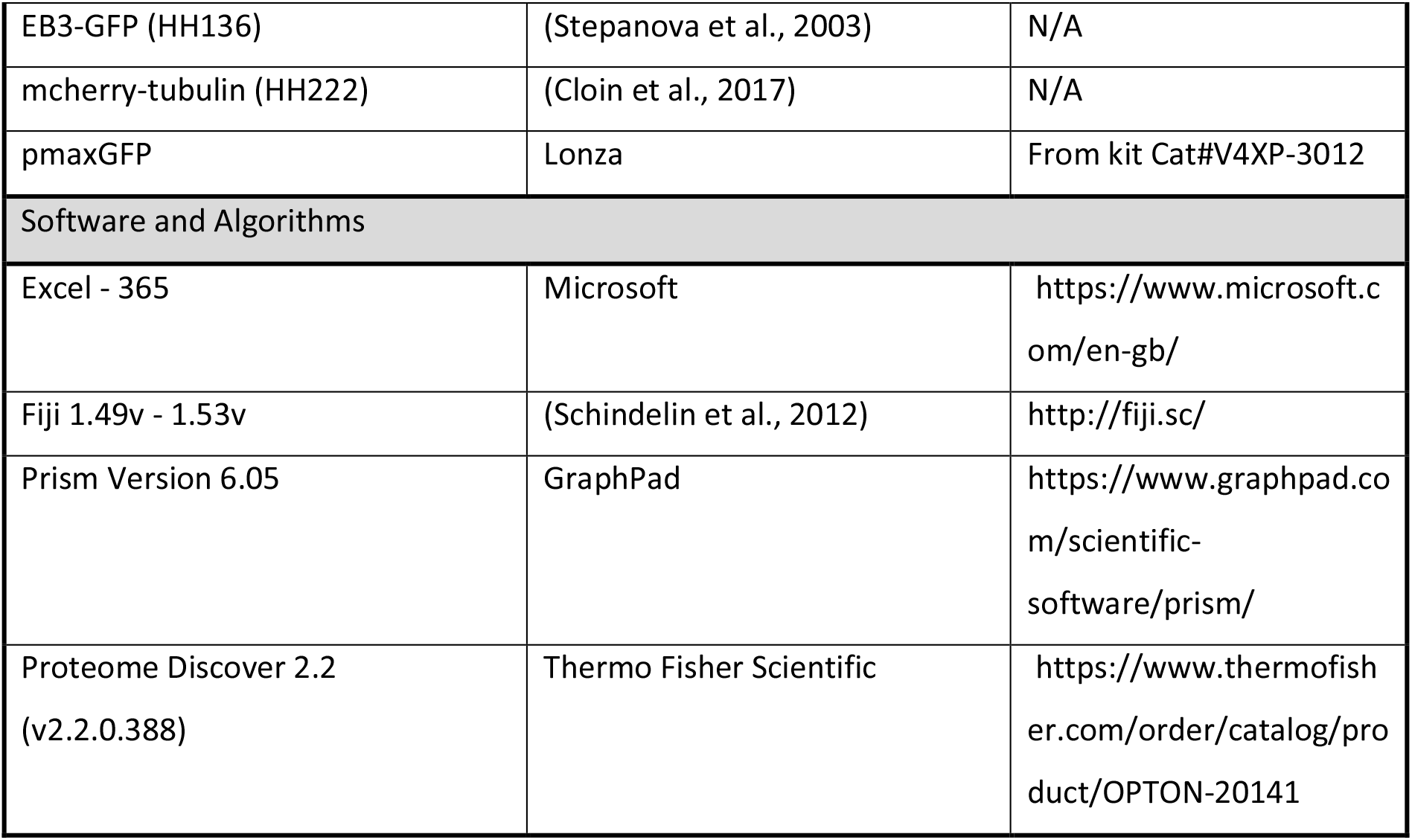
Crucial reagents and resources

### Cell Culture, transfections and treatments

HEK293T cells were grown in DMEM (Thermo Fisher #41966-029) +10 % FCS (Thermo Fisher #10270106) + 1x penicillin-streptomycin (Thermo Fisher #15140122) at 37 °C in 5 % CO_2_, 95 % humidity atmosphere and split twice per week. Cells for immunochemistry were plated on poly-L-lysin (Sigma P2636) coated glass coverslips, for biochemistry directly in 6-well plates. HEK293T cells were authenticated and tested for contamination by the supplier and used within 35 passages. Cells were transfected using maxPEI or lipofectamine 2000 (for siRNA experiments) using the manufacturers’ protocols 1 day after splitting cells and fixed or harvested after 24 h. For siRNA experiments, 15 pmol siRNA was used in combination with 0.6 μg DNA per well in a 12-well plate (1 ml volume). Only 0.3 μg pmaxGFP as a cell fill was used. Amounts were accordingly adjusted for use of 6-well plates. Treatments of HEK293T cells were performed so that all groups were fixed/harvested at the same time. The treatment details are: DMSO (1 h 1:1000), CPI (24 h 50 μg/ml), parthenolide (100 nM 24 h), calpeptin (60 μM 30 min), taxol (10 nM 30 min), calpastatin (20 μM 1h), PD151746 (260 nM 1 h).

### Immunocytochemistry (ICC)

ICC on HEK293T cells was performed as described previously (Bär et al., 2016). Briefly, cells were fixed with 4 % PFA/4 % sucrose, washed in PBS, permeabilized in 0.2 % Triton X-100 in PBS and blocked in blocking buffer (BB, 10 % horse serum, 1 % Triton X-100 in PBS). Primary antibodies were diluted in BB (anti-detyr-α-tubulin 1:400, anti-α-tubulin 1:600) and coverslips incubated over night at 4 °C. After additional washing with PBS, corresponding fluorescently labelled secondary antibodies (all 1:500 diluted in BB) were applied for 1-2 h at room temperature, before final washing with PBS and mounting of coverslips on objective slides with mowiol.

### Cell lysates and Western Blots

Cell homogenates/extracts of HEK293T cells were prepared as follows: Cells were shortly washed with warm PBS, harvested in Tris-buffered saline (TBS, 20 mM Tris, 150 mM NaCl, pH 7.4) + 1 % Triton X-100 + protease inhibitor cocktail (PI)). Hot 4x SDS sample buffer (250mM Tris-HCl, pH 6.8, 8 % (w/v) SDS, 40 % (v/v) Glycerol, 20 % (v/v) β-mercaptoethanol, 0.008 % bromophenol blue, pH 6.8) was added directly and samples boiled for 5 min. Equal amounts of samples were separated on a 10 % polyacrylamide gel and transferred to a PVDF membrane.

The membranes were blocked for 1 h in 5 % milk in Tris-buffered saline with 0.1 % Tween-20 at room temperature and incubated overnight at 4 °C with primary antibodies in TBS with 0.02 % NaN_3_ (anti-detyr-α-tubulin 1:1000, anti-β-actin 1:5000, anti-tyr-α-tubulin 1:1000, anti-calpain1 1:200, anti-calpain 2 1:200, vasohibin 1 1:100, anti-GAPDH 1:200. After washing in TBS + 0.1 % Tween and TBS, the membranes were incubated with secondary antibodies in 5 % milk in TBS + 0.1 % Tween for 1-2 h at room temperature, washed, and signals detected using ECL solution on an INTAS ChemoCam Imager (Intas Science Imaging). Detyr-α-tubulin, Tyr- α-tubulin and total α-tubulin were detected on different membranes, and the loading (actin, GAPDH) was detected on identical membranes and used for normalization. The Fiji tool “Analyze gels” was used to quantify band intensities.

### Microscopy

#### Widefield

Widefield imaging was performed at a Nikon Eclipse Ti-E microscope controlled by VisiView software and equipped with standard GFP, RFP, and Cy5 filters. Illumination was achieved by an LED light source. Use of a 100x (*Nikon*, ApoTIRF 1.49 oil), 60x 1.4NA (*Nikon*, CFI Plan Apo Lambda Oil) or 40x 1.3NA (*Nikon*, CFI Plan Fluor Oil) objective for imaging for transfected HEK293T or neurons, yielded pixel sizes of 65 nm, 108 nm or 162.5 nm. Images were taken at 16 bit depth and 2048 x 2048 pixel on an Orca flash 4.0LT CMOS camera (*Hamamatsu*). In part, several images on different z-positions were taken to cover the complete cells and maximum projections calculated for representation.

#### Confocal microscopy

Confocal imaging was performed at a Leica TCS SP5 confocal microscope (*Leica microsystems, Mannheim, Germany*), controlled by Leica Application Suite Advanced Fluorescence software. Samples were imaged using a 63x oil objective (*Leica,* 63x HCX PL APO Lbd. Bl. Oil/1.40 oil). Fluorophores were excited with Multi-Argon 488 nm, Diode Pumped Solid State 561 nm, HeNe 633 nm lasers and signals detected using HyD detectors. Pixel depth of 8 bit and frame averaging of 2 was used. Dimensions were set to 1024 x 1024 pixels and zoom set so that the resulting pixel size is 120 nm. Images were acquired at 400 Hz and as z-stacks with a step size of 0.29 μm.

#### Live imaging of EB3-GFP comets

Imaging of EB3 comets was performed 24h after transfection of HEK293T cells with EB3-GFP and tubulin-mCherry in combination with the different siRNAs. A Nikon Eclipse Ti-E controlled by VisiView software was used. Illumination was done by 488 nm excitation laser excitation controlled by a Targeted Laser Illuminator 2 (iLas2, *Gataca Systems*) in total internal reflection fluorescence (TIRF) mode/oblique illumination. Emission was collected through a quad band filter (*Chroma*, ZET 405/488/561/647m) and emission filters (*Chroma*, ET525/50m, ET595/50m, ET700/75m) on an Orca flash 4.0LT CMOS camera (*Hamamatsu*). The use of a 100x TIRF objective (*Nikon*, ApoTIRF 100x/1.49 oil) resulted in a pixel size of 65 nm. Images were acquired at 2-3 Hz with 200-300 ms illumination.

### Pulldown and *in vitro* calpain cleavage of flag-vash1-GFP + SVBP-myc for mass spectrometry analysis of cleavage sites

HEK293T cells were transfected with flag-vash1-GFP + SVBP-myc construct using maxPEI. After 24 h cells were washed with cold TBS (20 mM Tris, 150 mM NaCl, pH 8.0), and harvested in extraction buffer (20 mM Tris, 150 mM NaCl, 0.5 % Triton X-100, 2x PI, pH 8.0). Extraction was performed for 30 min on ice, followed by centrifugation at 14000 x g at 4 °C and supernatant used. Myc-Trap agarose beads A (Chromotek, yta-2 Myc Trap_A) were prepared by 3 rounds of washing in extraction buffer and centrifugation at 1000 x g for 5min at 4 °C. Beads were incubated with HEK extracts at 4 °C overnight at slow rotation. Unbound fraction was removed by centrifugation at 1000 x g for 5 min at 4 °C. Beads were washed 3 times in extraction buffer and equally split into 2 samples for on bead calpain digestion and control. Beads were washed 3 times with TBS to remove protease inhibitors and washed once with TBS + 2 mM CaCl_2_. 50 μl TBS + CaCl_2_ (all component 1.2x concentrated) was added to both samples. In vitro calpain cleavage was achieved by adding 10 μl calpain (>20 U, resulting in 1x concentration of TBS) and incubation at 30 °C for 5 min. The reaction was stopped by addition of 20 μl hot 4x SDS buffer, and the sample was boiled for 5 min. The control sample was treated equally except that 10 μl H_2_O were added instead of calpain. Samples were separated on a commercial 4-12 % bis tris plus SDS gel (Thermo Fisher #NW04120BOX).

### Mass spectrometry

Gel bands were cut into 2 mm^3^, destained using 100 mM ammonium bicarbonate (ABC), 30% acetonitrile (ACN) in 50 mM ABC buffer followed by dehydration with ACN and then dried using vacuum centrifugation. Samples were reduced with dithiothreitol (10 mM) in TEAB buffer (pH 8.5) for 30 min at 60°C and alkylated with iodoacetamide (55 mM) at room temperature for 15 min.

The gel pieces were washed with 30 % ACN, in HEPES buffer (25 mM) and then dehydrated with ACN. After drying the amino groups (N-terminus and lysine residues) were reductively dimethylated using 40 mM formaldehyde, 20 mM sodium cyanoborohydride in HEPES buffer (250 mM, pH 7) at 37°C for 16 h. The reaction was quenched using 0.9 M of Tris buffer (pH 6.8) for 3 hr followed by multiple washing steps, i.e., 50 mM ABC in 30 % ACN (15 min), 100% ACN (15 min), drying in a vacuum centrifuge (10 min). After quenching the gel bands were incubated with enzyme either trypsin (50 ng) or chymotrypsin (100 ng) in the presence of 50% H_2_^18^O (heavy water). Trypsin samples were incubated with 25 mM ABC, while bands digested chymotrypsin also included 2 mM CaCl_2_ in the digestion buffer. Samples were incubated overnight a 37°C and after digestion the reaction was quenched using 10 % formic acid (FA). The peptides from the gel bands were extracted with 50% ACN in 1% FA and 100% ACN using sonication. Pooled extracts were dried down and the samples resuspended in 3% ACN, 0.1% trifluoroacetic acid (TFA) prior to LC-MS.

In-gel digested samples were analyzed on a Dionex Ultimate 3000 nano-UHPLC coupled to a Q Exactive mass spectrometer (Thermo Scientific, Bremen). The samples were washed on a trap column (Acclaim Pepmap 100 C18, 5 mm × 300 μm, 5 μm, 100 Å, Dionex) for 4 min with 3 % ACN/0.1 % TFA at a flow rate of 30 μl/min prior to peptide separation using an Acclaim PepMap 100 C-18 analytical column (50 cm × 75 μm, 2 μm, 100 Å, Dionex). A flow rate of 300 nL/min using eluent A (0.05 % FA) and eluent B (80 % ACN/0.04 % FA) was used for gradient separation. Spray voltage applied on a metal-coated PicoTip emitter (10 μm tip size, New Objective, Woburn, Massachusetts, US) was 1.7 kV, with a source temperature of 250 °C. Full scan MS spectra were acquired between 300 and 2,000 *m/z* at a resolution of 70,000 at *m/z* 400. The ten most intense precursors with charge states greater than 2+ were selected with an isolation window of 2.1 *m/z* and fragmented by HCD with normalized collision energies of 27 at a resolution of 17,500. Lock mass (445.120025) and dynamic exclusion (15 s) were enabled.

The MS raw files were processed by Proteome Discover 2.2 (Thermo, version 2.2.0.388) and MS/MS spectra were searched using the Sequest HT algorithm against a database containing common contaminants (45 sequences), the canonical human database and recombinant flag-vash1-GFP-his. For trypsin, the enzyme specificity was set to semi-Arg-C specificity with two missed cleavages allowed. For samples incubated with chymotrypsin, no enzyme specificity was used. An MS1 tolerance of 10 ppm and a MS2 tolerance of 0.02 Da was implemented. Oxidation (15.995 Da) of methionine residues, dimethylation (28.031 Da) on the peptide N-terminus, and heavy water (2.004 Da) on the peptide C-terminus were set as a variable modification. Carbamidomethylation (57.02146 Da) on cysteine residues along with dimethylation on lysine residues were set as a static modification. Minimal peptide length was set to 6 amino acids and the peptide false discovery rate (FDR) was set to 1 %. Peptide peak intensities were calculated using the Minora. Peak intensities were only used if they were identified as high-confident.

Abundances of all the N-termini that were identified with high confidence for bands 2,3,4 and 1 (control) for both chymotrypsin (Supplemental Table 1) and trypsin (Supplemental Table 1) digestions. Samples were analyzed from lowest molecular weight to highest molecular weight, with control measured last. The same volume of digested peptide extract was loaded for LC-MS for all samples. All peptides shown have their N-terminus and their lysine residues dimethylated. The large number of N-termini identified in the control sample may be due to degradation of vash1 or insufficient quenching of labeling reagents prior to enzymatic digestion. N-termini were only considered to be calpain cleavage events if they were identified in both the chymotrypsin and trypsin experiments and were also not observed in the calpain control. Peptide was also identified in control sample, but with very low intensity and likely to be a result of carryover. Most probably calpain cleavage sites, based on intensity and number of PSMs, are highlighted.

### Purification of twin-strepTag-vash1-GFP for *in vitro* MT assays

HEK293T cells were transfected at 60-70 % confluency using maxPEI with twin-strep-tag-vash1-GFP IRES SVBP construct 24 h before purification. Cells were washed once with ice-cold TBS and harvested in ice-cold TBS subsequently. The cells were centrifuged at 1000 x g for 3 min at 4 °C and resuspended in extraction buffer (20 mM Tris, 300 mM NaCl, 1 % Triton X-100, 5 mM MgCl_2_, 1 x cOmplete, EDTA-free Protease Inhibitor Cocktail (Sigma), pH 8.0) and incubated for 30 min on ice. The cell lysate was centrifuged at 14000 x g for 15 min at 4 °C. The supernatant was collected and incubated with Strep-Tactin Sepharose High Performance beads (GE Healthcare) for 1 h at 4 °C on slow rotation. The beads were centrifuged at 1000 x g for 1 min at 4 °C and the cell lysate was taken off. Beads were washed 3 times with washing buffer (20 mM Tris, 150 mM NaCl, 0.5 % Triton X-100, 2 mM EGTA, 1 x cOmplete, EDTA-free Protease Inhibitor Cocktail (Sigma), pH 8.0). Elution was performed 3 times for 10 min incubation in elution buffer (100 mM Tris, 150 mM NaCl, 1 mM EDTA, 50 mM D-Biotin, 1 mM DTT) followed by centrifugation at 1000 x g for 1 min.

The purified protein was stored on ice until further use. Purification was controlled by adding 5 μL of 2 x SDS buffer to 5 μL of first elution, boiling the sample for 5 min and running SDS-PAGE with BSA standard on a 10 % SDS-polyacrylamide gel. The gel was stained with Coomassie Brilliant Blue R250 (Roth) for 30 min at room temperature and destained with MilliQ-water over night at room temperature.

### *In vitro* cleavage of purified twin-strepTag-vash1-GFP by calpain 1

μM purified twin-strepTag-vash1-GFP was incubated with 5 μL calpain1 (>10 U) in 100 mM TBS with 2 mM CaCl_2_ for 5 min at 37°C. Afterwards, the samples were put on ice immediately and used straight away.

The Calpain1 cleavage of twin-StrepTag-vash1-GFP was analyzed by SDS-PAGE with 12 % SDS-polyacrylamide gel and followed by western blot (procedure: see above). Blocking was done for 30 min with 5 % milk powder in TBS-T. Primary antibodies used were anti-calpain1 (1:100 in TBS with 0.02 % NaN_3_), anti-vasohibin 1 (1:100 in TBS with 0.02 % NaN_3_). Secondary antibody used was anti-mouse-HRP (1:20000 in TBS-T). The blots were developed using ECL solution on an INTAS ChemoCam Imager.

### *In vitro* MT detyrosination assay

rounds of MT polymerization and cold-induced depolymerization were performed to enrich for tyrosinated-tubulin. A mix of 20 μM of porcine brain tubulin protein (Cytoskeleton/Tebu-Bio) with 1 mM GMPCPP (Jena Bioscience) in PEM80 (80 mM Pipes, 1 mM EGTA, 4 mM MgCl_2_) was made on ice and subsequently incubated at 37 °C for 30 min for production of microtubule seeds. The sample was centrifuged at 120000 x g for 5 min at 25 °C to remove polymerized MTs from solution. The supernatant was discarded, and the pellet was resuspended in PEM80 to about 20 μM tubulin assuming 80 % recovery. The resuspended tubulin was incubated for 20 min on ice to depolymerize the MTs. Then GMPCPP was added to a concentration of 1 mM and incubated 5 min more on ice and subsequently, the process was repeated.

After the second round of MT polymerization and depolymerization, the sample was incubated for 30 min at 37 °C to obtain MTs. 0.1 μM of purified twin-StrepTag-vash1-GFP, 0.1 μM *in vitro* calpain1 cleaved twin-StrepTag-vash1-GFP (see above) or elution buffer (see above) without protein as negative control were added to MTs and incubated for 5, 10, 20 or 30 min at 37 °C. The reaction was stopped by addition of hot 4 x SDS buffer, and samples were boiled for 5 min.

Equal amounts of the samples were analyzed by SDS-PAGE with 12 % SDS-polyacrylamide gel, followed by western blot (procedure: see above). Blocking was done for 30 min with 5 % milk powder in TBS-T. Primary antibodies used were anti-detyr-α-tubulin (1:1000 in 5 % milk powder in TBS-T), anti-tyr-α-tubulin (1:1000 in 5 % milk powder in TBS-T) and anti- α-tubulin (1:2000 in TBS with 0.02 % NaN_3_). Secondary antibodies used were anti-rabbit-HRP (1:20000 in TBS-T) and anti-mouse-HRP (1:20000 in TBS-T). The blots were developed using ECL solution on an INTAS ChemoCam Device. Band intensity integration areas were analyzed using the “Analyze gels” tool of Fiji. The ratio tyr-α-tubulin/anti-α-tubulin was calculated and normalized to the corresponding control.

### Data analysis

#### Comets tracks and counts

2 min of live imaging was analyzed after applying a background subtraction. EB3 comets were detected and analyzed using the Fiji (Schindelin et al., 2012) plugin “TrackMate” version v5 (Tinevez et al., 2017) using the following settings: Particle detection with LoG detection, initial particle size of approx 0.4-0.5 μm, threshold set individually per video., no median filter applied, with subpixel localization and without quality filters for the detection of spots themselves. For tracking the simple LAP tracker was used with the following settings: 0.2-0.5 μm linking max distance, 0.5-0.8 μm gap-closing distance and 0-6 gap closing max frame gap. Tracks below 5 s tracking length were excluded to remove very short and unreliably tracked comets. This may result in insensitivity to changes in very short tracks but allows the proper quantification of EB track speeds. To estimate the number of EB3 comets, the number of tracks containing the first frame was counted.

#### ICC quantification

Detyr-α-tubulin/α-tubulin ratios in immunostainings were analyzed as follows: Cells were outlined by using the total tubulin channel, that was after strong smoothing and despeckling, semi-automatically thresholded and manually corrected to exclude dividing cells, dense cell clusters and parts of cells on the edge of the field of view. Afterwards, the GFP channel was smoothed and despeckled and used to identify transfected cells in the same way, and cell outlines were corrected with the help of the total tubulin channel. Subtraction of the total cell outline and the transfected cell outline was calculated to obtain the outline of non-transfected cells. Total tubulin and detyr-tubulin average intensity values within the transfected and untransfected cells were measured in the original images. For each image, the detyr-α-tubulin/α-tubulin ratio of transfected and untransfected cells was calculated and afterwards intensities of transfected cells normalized to untransfected cells. Data from several 3 independent experiments were normalized to control, before pooling data. An experimenter blinded to the experimental groups performed all analyses.

### Statistical analysis and image representation

Statistical analysis was performed using Prism version 6.07 (GraphPad). Data are represented as mean ± SEM. Data were tested for normal distribution (D’Agostino & Pearson omnibus normality test) and statistical test chosen accordingly. Individual channels of multi-color micrographs were contrasted for better representation, with identical settings within 1 experiment. All analysis was performed on raw images.

## Acknowledgements

We would like to thank the CNI Imaging facility of the Leibniz Institute for Neurobiology, Magdeburg, for access to their Leica SP5 confocal microscope, Daniela Hacker for help with initial antibody characterizations, Marie-Jo Moutin for kindly providing the flag-full-length-vash1-sfGFP-his IRES SVBP-myc and the vash knockdown constructs, as well as Amol Aher and Anna Akhmanova for kindly providing the twinStrep-mCherry empty vector.

## Data availability

N/A

## References

Adamopoulos, A., Landskron, L., Heidebrecht, T., Tsakou, F., Bleijerveld, O. B., Altelaar, M., Nieuwenhuis, J., Celie, P. H. N., Brummelkamp, T. R. and Perrakis, A. (2019). Crystal structure of the tubulin tyrosine carboxypeptidase complex VASH1–SVBP. Nature Structural & Molecular Biology 26, 567–570.

Aillaud, C., Bosc, C., Peris, L., Bosson, A., Heemeryck, P., Van Dijk, J., Le Friec, J., Boulan, B., Vossier, F., Sanman, L. E. et al. (2017). Vasohibins/SVBP are tubulin carboxypeptidases (TCPs) that regulate neuron differentiation. Science 358, 1448–1453.

Amini, M., Ma, C.-l., Farazifard, R., Zhu, G., Zhang, Y., Vanderluit, J., Zoltewicz, J. S., Hage, F., Savitt, J. M., Lagace, D. C. et al. (2013). Conditional Disruption of Calpain in the CNS Alters Dendrite Morphology, Impairs LTP, and Promotes Neuronal Survival following Injury. The Journal of Neuroscience 33, 5773–5784.

Bär, J., Kobler, O., van Bommel, B. and Mikhaylova, M. (2016). Periodic F-actin structures shape the neck of dendritic spines. Scientific Reports 6, 37136.

Barisic, M., Silva e Sousa, R., Tripathy, S. K., Magiera, M. M., Zaytsev, A. V., Pereira, A. L., Janke, C., Grishchuk, E. L. and Maiato, H. (2015). Microtubule detyrosination guides chromosomes during mitosis. Science 348, 799–803.

Chen, C. Y., Caporizzo, M. A., Bedi, K., Vite, A., Bogush, A. I., Robison, P., Heffler, J. G., Salomon, A. K., Kelly, N. A., Babu, A. et al. (2018). Suppression of detyrosinated microtubules improves cardiomyocyte function in human heart failure. Nat Med 24, 1225–1233.

Cloin, B. M. C., De Zitter, E., Salas, D., Gielen, V., Folkers, G. E., Mikhaylova, M., Bergeler, M., Krajnik, B., Harvey, J., Hoogenraad, C. C. et al. (2017). Efficient switching of mCherry fluorescence using chemical caging. Proceedings of the National Academy of Sciences 114, 7013–7018.

Deng, J., Zhang, G., Huang, F.-K. and Neubert, T. A. (2015). Identification of Protein N-Termini Using TMPP or Dimethyl Labeling and Mass Spectrometry. In Proteomic Profiling: Methods and Protocols, (ed. A. Posch), pp. 249–258. New York, NY: Springer New York.

Desai, A. and Mitchison, T. J. (1997). Microtubule polymerization dynamics. Annu Rev Cell Dev Biol 13, 83–117.

Dimitrov, A., Quesnoit, M., Moutel, S., Cantaloube, I., Poüs, C. and Perez, F. (2008). Detection of GTP-Tubulin Conformation in Vivo Reveals a Role for GTP Remnants in Microtubule Rescues. Science 322, 1353–1356.

Dubey, J., Ratnakaran, N. and Koushika, S. (2015). Neurodegeneration and microtubule dynamics: death by a thousand cuts. Front Cell Neurosci 9.

Garnham, C. P. and Roll-Mecak, A. (2012). The chemical complexity of cellular microtubules: Tubulin post-translational modification enzymes and their roles in tuning microtubule functions. Cytoskeleton 69, 442–463.

Heinz, L. S., Muhs, S., Schiewek, J., Grüb, S., Nalaskowski, M., Lin, Y.-N., Wikman, H., Oliveira-Ferrer, L., Lange, T., Wellbrock, J. et al. (2017). Strong fascin expression promotes metastasis independent of its F-actin bundling activity. Oncotarget 8.

Honda, S., Marumoto, T., Hirota, T., Nitta, M., Arima, Y., Ogawa, M. and Saya, H. (2004). Activation of m-calpain is required for chromosome alignment on the metaphase plate during mitosis. J Biol Chem 279, 10615–23.

Iqbal, Z., Tawamie, H., Ba, W., Reis, A., Halak, B. A., Sticht, H., Uebe, S., Kasri, N. N., Riazuddin, S., van Bokhoven, H. et al. (2019). Loss of function of SVBP leads to autosomal recessive intellectual disability, microcephaly, ataxia, and hypotonia. Genet Med 21, 1790–1796.

Janke, C. (2014). The tubulin code: Molecular components, readout mechanisms, and functions. Journal of Cell Biology 206, 461–472.

Jurriens, D., van Batenburg, V., Katrukha, E. A. and Kapitein, L. C. (2021). Mapping the neuronal cytoskeleton using expansion microscopy. Methods Cell Biol 161, 105–124.

Kadonosono, T., Yimchuen, W., Tsubaki, T., Shiozawa, T., Suzuki, Y., Kuchimaru, T., Sato, Y. and Kizaka-Kondoh, S. (2017). Domain architecture of vasohibins required for their chaperone-dependent unconventional extracellular release. Protein Science 26, 452–463.

Kaul, N., Soppina, V. and Verhey, K. J. (2014). Effects of α-tubulin K40 acetylation and detyrosination on kinesin-1 motility in a purified system. Biophys J 106, 2636–43.

Li, F., Hu, Y., Qi, S., Luo, X. and Yu, H. (2019). Structural basis of tubulin detyrosination by vasohibins. Nature Structural & Molecular Biology 26, 583–591.

Liao, S., Rajendraprasad, G., Wang, N., Eibes, S., Gao, J., Yu, H., Wu, G., Tu, X., Huang, H., Barisic, M. et al. (2019). Molecular basis of vasohibins-mediated detyrosination and its impact on spindle function and mitosis. Cell Research 29, 533–547.

Mingorance-Le Meur, A. and O’Connor, T. P. (2009). Neurite consolidation is an active process requiring constant repression of protrusive activity. Embo j 28, 248–260.

Moldoveanu, T., Hosfield, C. M., Lim, D., Elce, J. S., Jia, Z. and Davies, P. L. (2002). A Ca(2+) switch aligns the active site of calpain. Cell 108, 649–60.

Nieuwenhuis, J., Adamopoulos, A., Bleijerveld, O. B., Mazouzi, A., Stickel, E., Celie, P., Altelaar, M., Knipscheer, P., Perrakis, A., Blomen, V. A. et al. (2017). Vasohibins encode tubulin detyrosinating activity. Science 358, 1453–1456.

Ono, Y., Saido, T. C. and Sorimachi, H. (2016). Calpain research for drug discovery: challenges and potential. Nature Reviews Drug Discovery 15, 854–876.

Pagnamenta, A. T., Heemeryck, P., Martin, H. C., Bosc, C., Peris, L., Uszynski, I., Gory-Faure, S., Couly, S., Deshpande, C., Siddiqui, A. et al. (2019). Defective tubulin detyrosination causes structural brain abnormalities with cognitive deficiency in humans and mice. Hum Mol Genet 28, 3391–3405.

Rawlings, N. D., Barrett, A. J., Thomas, P. D., Huang, X., Bateman, A. and Finn, R. D. (2017). The MEROPS database of proteolytic enzymes, their substrates and inhibitors in 2017 and a comparison with peptidases in the PANTHER database. Nucleic Acids Research 46, D624–D632.

Redeker, V. (2010). Chapter 6 - Mass Spectrometry Analysis of C-Terminal Posttranslational Modifications of Tubulins. In Methods Cell Biol, vol. 95 (eds L. Wilson and J. J. Correia), pp. 77–103: Academic Press.

Saito, M., Suzuki, Y., Yano, S., Miyazaki, T. and Sato, Y. (2016). Proteolytic inactivation of anti-angiogenic vasohibin-1 by cancer cells. The Journal of Biochemistry 160, 227–232.

Schindelin, J., Arganda-Carreras, I., Frise, E., Kaynig, V., Longair, M., Pietzsch, T., Preibisch, S., Rueden, C., Saalfeld, S., Schmid, B. et al. (2012). Fiji: an open-source platform for biological-image analysis. Nature Methods 9, 676–682.

Schnölzer, M., Jedrzejewski, P. and Lehmann, W. D. (1996). Protease-catalyzed incorporation of 18O into peptide fragments and its application for protein sequencing by electrospray and matrix-assisted laser desorption/ionization mass spectrometry. ELECTROPHORESIS 17, 945–953.

Seidah, N. G. and Chrétien, M. (1999). Proprotein and prohormone convertases: a family of subtilases generating diverse bioactive polypeptides. Brain Research 848, 45–62.

Shinkai-Ouchi, F., Koyama, S., Ono, Y., Hata, S., Ojima, K., Shindo, M., duVerle, D., Ueno, M., Kitamura, F., Doi, N. et al. (2016). Predictions of Cleavability of Calpain Proteolysis by Quantitative Structure-Activity Relationship Analysis Using Newly Determined Cleavage Sites and Catalytic Efficiencies of an Oligopeptide Array. Mol Cell Proteomics 15, 1262–1280.

Sirajuddin, M., Rice, L. M. and Vale, R. D. (2014). Regulation of microtubule motors by tubulin isotypes and post-translational modifications. Nat Cell Biol 16, 335–344.

Sonoda, H., Ohta, H., Watanabe, K., Yamashita, H., Kimura, H. and Sato, Y. (2006). Multiple processing forms and their biological activities of a novel angiogenesis inhibitor vasohibin. Biochem Biophys Res Commun 342, 640–646.

Sorimachi, H., Mamitsuka, H. and Ono, Y. (2012). Understanding the substrate specificity of conventional calpains. Biological Chemistry 393, 853–871.

Stepanova, T., Slemmer, J., Hoogenraad, C. C., Lansbergen, G., Dortland, B., De Zeeuw, C. I., Grosveld, F., van Cappellen, G., Akhmanova, A. and Galjart, N. (2003). Visualization of Microtubule Growth in Cultured Neurons via the Use of EB3-GFP (End-Binding Protein 3-Green Fluorescent Protein). The Journal of Neuroscience 23, 2655–2664.

Tas, R. P., Chazeau, A., Cloin, B. M. C., Lambers, M. L. A., Hoogenraad, C. C. and Kapitein, L. C. (2017). Differentiation between Oppositely Oriented Microtubules Controls Polarized Neuronal Transport. Neuron 96, 1264–1271.e5.

Théry, M. and Blanchoin, L. (2021). Microtubule self-repair. Curr Opin Cell Biol 68, 144–154.

Tinevez, J.-Y., Perry, N., Schindelin, J., Hoopes, G. M., Reynolds, G. D., Laplantine, E., Bednarek, S. Y., Shorte, S. L. and Eliceiri, K. W. (2017). TrackMate: An open and extensible platform for single-particle tracking. Methods 115, 80–90.

Uhlén, M., Fagerberg, L., Hallström, B. M., Lindskog, C., Oksvold, P., Mardinoglu, A., Sivertsson, Å., Kampf, C., Sjöstedt, E., Asplund, A. et al. (2015). Tissue-based map of the human proteome. Science 347, 1260419.

van der Laan, S., Lévêque, M. F., Marcellin, G., Vezenkov, L., Lannay, Y., Dubra, G., Bompard, G., Ovejero, S., Urbach, S., Burgess, A. et al. (2019). Evolutionary Divergence of Enzymatic Mechanisms for Tubulin Detyrosination. Cell Reports 29, 4159–4171.e6.

Wang, N., Bosc, C., Ryul Choi, S., Boulan, B., Peris, L., Olieric, N., Bao, H., Krichen, F., Chen, L., Andrieux, A. et al. (2019). Structural basis of tubulin detyrosination by the vasohibin–SVBP enzyme complex. Nature Structural & Molecular Biology 26, 571–582.

Yasuda, K., Clatterbuck-Soper, S. F., Jackrel, M. E., Shorter, J. and Mili, S. (2017). FUS inclusions disrupt RNA localization by sequestering kinesin-1 and inhibiting microtubule detyrosination. Journal of Cell Biology 216, 1015–1034.

Zadran, S., Bi, X. and Baudry, M. (2010). Regulation of Calpain-2 in Neurons: Implications for Synaptic Plasticity. Mol Neurobiol 42, 143–150.

